# A Gm6AG-binding protein from *Vibrio cholerae*

**DOI:** 10.1101/2025.05.22.655678

**Authors:** Yue Cao, Jialu Xu, Xueling Lu, Fengtao Huang, Weihua Chen, Xionglue Wang, Bin Zhu

## Abstract

Most known modification-dependent restriction endonucleases target 5-methylcytosine, only a few N6-methyladenine (6mA)-dependent restriction endonucleases have been well-characterized, and the majority of them recognize the G6mATC motif (e.g., DpnI, HHPV4I). Here, we report the identification of a novel 6mA-dependent DNA-binding protein from *Vibrio cholerae*, VchI, which specifically recognizes the G6mAG motif. VchI contains a winged helix (wH) domain that is homologous to the wH domain in DpnI. However, several key residues involved in 6mA recognition differ between VchI and DpnI, which may contribute to the discrepancy in their recognition specificities. These findings advance our understanding of prokaryotic 6mA modification diversity and the 6mA recognition mechanism of the wH domain, while simultaneously providing an innovative tool for epigenetic research.

## Introduction

DNA methylation represents a fundamental epigenetic modification (1,2). In prokaryotes, N6-methyladenine (6mA) is the predominant DNA methylation form, which plays pivotal roles in bacteriophage defense, DNA replication regulation, and cell cycle progression (3–5). Modification-dependent restriction endonucleases (MDREs), capable of recognizing and cleaving specifically modified DNA, serve as indispensable tools for epigenetic research (6–8). Most known MDREs target 5-methylcytosine (5mC) modification (9–15), while only a small number of 6mA-dependent restriction endonucleases have been well characterized (e.g., DpnI, HHPV4I) (16–18), which exhibit exclusive specificity for the G6mATC motif. This has led to a shortage of tool enzymes for studying 6mA modifications in epigenetic research. To address this limitation, we systematically explored natural sources to discover novel 6mA-dependent restriction endonucleases or 6mA-recognition proteins. Our investigation led to the identification of VchI, a novel 6mA-dependent DNA-binding protein from *Vibrio cholerae*, which specifically recognizes the G6mAG motif.

## Materials and methods

### Plasmid construction

The codon-optimized VchI gene (Protein ID: WP_057555222.1; GenBank ID: CP046743.1, CP013308.1, AP014525.1, CP016325.1, CP129422.1, CP053811.1, CP053814.1) was ordered from GenScript (Nanjing, China) and cloned into a pQE80L vector with an N-terminal 6×His tag.

### Protein expression and purification

Vch? was expressed in *E. coli* Trans110 (Dam- and Dcm-). The protein-expression plasmid of Vch? was transformed into *E. coli* Trans110. The transformants were inoculated in LB medium and grown overnight at 37°C. The cultures were transferred into fresh LB medium at a ratio of approximately 1:100. When the OD600 reached approximately 1.0, IPTG (isopropyl-β-D-thiogalactopyranoside) was added at 0.5 mM, and the cultures were incubated at 25°C for 5 h. Then, the cells were harvested and resuspended in lysis buffer (20 mM Tris-HCl, pH 7.9, 300 mM NaCl, 20 mM imidazole) and lysed by sonication on ice. The recombinant proteins were purified using Ni-NTA resin (Qiagen) and a gravity column. The Ni-NTA resin was washed (using lysis buffer supplemented to 50 mM imidazole) and eluted (using lysis buffer supplemented to 200 mM imidazole), and the eluate was dialyzed twice at 4°C against a storage buffer (50 mM Tris-HCl, pH 7.9, 100 mM NaCl, 0.1 mM EDTA, 1 mM DTT, 0.1% TritonX-100, and 50% glycerol).

### Preparation of DNA substrates

The 300 bp substrate was amplified via a two-step PCR using PrimeSTAR Max DNA polymerase with the pBR322 plasmid as the template. Oligonucleotides (≤60 bp), including 6-FAM-labeled and unmodified DNA and RNA strands, were synthesized by GenScript Biotech (Nanjing, China). The dsDNA was carried out on a PCR instrument with a specific annealing program (1. 90°C, 1 min; 2. 90°C, 5 s, −0.1°C per cycle; 3. GOTO step 2, 650×; 4. 4°C, ∞.). The reaction mixture consisted of 10 mM Tris-HCl (pH 7.9), 50 mM KCl, 25 μM 6-FAM-labeled DNA oligonucleotide or 25 μM unlabeled DNA oligonucleotide.

Dam methyltransferase reactions were performed in 1× Dam methyltransferase buffer (50 mM Tris-HCl, 5 mM β-mercaptoethanol, 10 mM EDTA, pH 7.5) containing 80 μM S-adenosylmethionine (SAM) and 8 U of Dam methyltransferase. For EcoGII methyltransferase (M.EcoGII) assays, reactions were conducted in 1× rCutSmart buffer (50 mM potassium acetate, 20 mM Tris-acetate, 10 mM magnesium acetate, 100 μg/ml recombinant albumin, pH 7.9) supplemented with 160 μM SAM and 5 U of M.EcoGII. EcoRI methyltransferase (M.EcoRI) reactions utilized 1× rCutSmart buffer with 80 μM SAM and 40 U of enzyme. All methylation reactions (Dam, M.EcoGII, and M.EcoRI) were performed using 1 μg of substrate DNA at 37°C for 1 h. TaqI methyltransferase (M.TaqI) reactions were carried out under distinct conditions: 1× rCutSmart buffer with 80 μM SAM and 10 U of enzyme, with incubation at 65°C for 1 h using 1 μg of DNA substrate.

### Electrophoretic mobility shift assay (EMSA)

Binding reactions (10 μL final volume) containing 50 nM nonmethylated DNA or various amounts of methylated DNA and VchI were performed in 1× reaction buffer (20 mM Tris-HCl, pH 8.0, 200 mM NaCl) at 37°C for 5 min. Then, the reactions were mixed with 4 μL of 6× loading dye and examined by 1% TAE agarose gel or 6%, 12%, 20% nondenaturing PAGE.

### DNA cleavage assays

Standard cleavage reactions (10 μL final volume) containing 100ng nonmethylated DNA or various amounts of methylated DNA and VchI or mutants were performed in 1× cleavage reaction buffer (50 mM Tris-HCl, pH 7.9, 100 mM NaCl, 5mM Mn^2+^) at 37°C for 1h. Any variations in reaction conditions are indicated in the figure legends. Then, the reactions were mixed with 4 μL of 6× loading dye and examined by 1% TAE agarose gel or 6%, 12%, 20% nondenaturing PAGE. The gel was stained with ethidium bromide and DNA or RNA was visualized with a UVsolo Touch system (Analytik Jena, Germany). The detection method varies depending on the cleavage substrate. For fluorescence-labeled dsDNA and ssRNA reaction substrates, the 6-FAM-labeled substrates should be observed using the Cy2 channel (Ex470BL, Em525/30F) of the ChemiScope imaging system.

### AlphaFold structural prediction

The structure of VchI was predicted using AlphaFold 3. The FASTA-formatted sequence of the target protein was submitted to the official AlphaFold server (https://alphafoldserver.com/) with default template mode, disabled multimer prediction, and a maximum of 3 recycles set as parameters. Among the five optimal prediction models output by the system, we prioritized models with a global mean pLDDT > 90 and key functional domain pLDDT > 70, while also referring to the predicted aligned error (PAE) matrix to screen models with stable structural domains. The final selected prediction model was analyzed and visualized using PyMOL for detailed structural and functional characterization, including secondary structure element visualization (cartoon representation), key amino acid residue labeling (stick representation), and structural quality validation (Ramachandran plot analysis). This prediction method requires only amino acid sequence information to achieve three-dimensional structure prediction with near-experimental accuracy.

### Data processing and analysis

For protein sequence analysis, we first retrieved the amino acid sequences of target proteins from the NCBI database in FASTA format. Multiple sequence alignment was then performed using Clustal W, and the resulting alignment files were subsequently analyzed through the jalview online platform to visualize conserved regions and predict potential functional motifs.

## Results

### A wH domain-containing protein from *Vibrio cholerae* recognizes a specific 6mA motif beyond G6mATC, GA6mTTC, and TCG6mA

Previous studies have demonstrated that the winged helix (wH) domain plays a crucial role in 6mA recognition (19). Through a homology analysis of the wH domain of DpnI, we found a protein with an N-terminal wH domain and a predicted C-terminal HNH endonuclease domain from *Vibrio cholerae*, which may function as a 6mA-dependent restriction endonuclease (Figure 1A). We designated this protein as VchI. To study the VchI activity *in vitro*, we constructed the VchI expression plasmid and overexpressed the protein in *E. coli*. The recombinant VchI protein was then purified using a Ni-NTA column to achieve high purity (Figure 1B).

**Figure 1.**
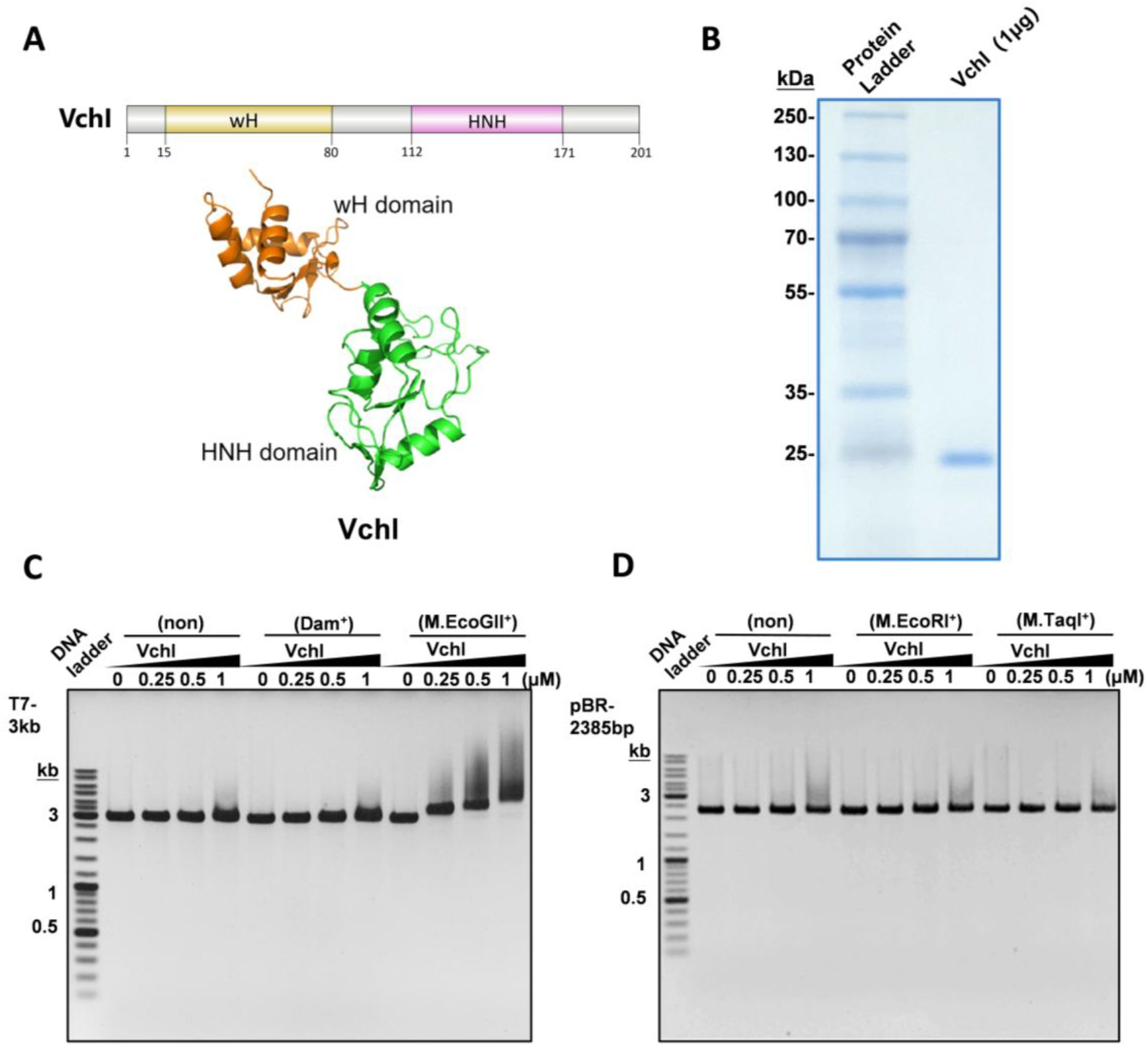
A wH domain-containing protein from Vibrio cholerae recognizes a specific 6mA motif beyond G6mATC, GA6mTTC, and TCG6mA. **(A)** Schematic diagram of the domain architectures and alphaFold structural predictions of VchI. **(B)**SDS-PAGE analysis of the purified VchI. **(C)** Binding of VchI to unmethylated, Dam-methylated, and M.EcoGII-methylated plasmids at varying concentrations. The substrate used in this experiment is a 3-kb fragment from the T7 phage genome. The specific primers are shown in Table S2. **(D)** Binding of VchI to DNA with different 6mA methylation patterns. Electrophoretic mobility shift assay (EMSA) was performed using unmethylated, M.EcoRI-methylated, and M.TaqI-methylated DNA substrates incubated with increasing concentrations of VchI. The substrate used in this experiment is pBR322 plasmid with a length of 2385bp.

To investigate whether VchI recognizes and cleaves DNA with 6mA modification, we first incubated VchI with a 3-kb DNA substrate modified with EcoGII methyltransferase (M. EcoGII), which modifies adenine residues in any sequence context. Intriguingly, though no cleavage was observed, VchI exhibited evident binding to this substrate (Figure 1C). However, when VchI was incubated with the DNA substrate containing G6mATC (modified with dam methytransferase), GA6mATTC (modified with M. EcoRI), TCG6mA (modified with M. TaqI), or no 6mA modification, neither cleavage nor binding was observed (Figure 1C, D). The results indicated that VchI recognizes a specific 6mA motif beyond G6mATC, GA6mTTC, and TCG6mA.

### Identification of the specific recognition motif of VchI

We amplified a 300-bp DNA fragment from the pBR322 plasmid and methylated the DNA with M. EcoGII *in vitro* (Figure 2A). The electrophoretic mobility shift assay (EMSA) showed dose-dependent binding of VchI to the 300-bp methylated DNA (Figure 2B), confirming the presence of its recognition motif in this substrate. By comparative analysis of VchI binding to seven overlapping 60-bp DNA segments covering the 300-bp DNA fragment (Figure 2A, C), we delineated three regions containing the recognition motif of VchI by subtracting sequences shared with non-binding fragments (60bp-1, 60bp-4, and 60bp-6) from binding-positive fragments (60bp-2, 60bp-3, 60bp-5, and 60bp-7) (Figure 2A). Notably, these three regions all contain at least one “GCGAG” sequence (Figure 2A, marked with green), suggesting that GCG6mAG may represent the recognition motif of VchI. To validate this hypothesis, we generated the variants of the 60bp-2 and 60bp-7 substrates by replacing the “GCGAG” sequences with “TTTTT” (designated as 60bp-2-V and 60bp-7-V) (Figure 3A). Although EMSA revealed no binding of VchI to 60bp-2-V, 60bp-7-V exhibited significant binding (Figure 3B), suggesting that while GCG6mAG represents a recognition motif for VchI, 60bp-7-V contains at least one additional VchI recognition motif in addition to GCG6mAG. Further sequence analysis of 60bp-7-V revealed the presence of a “CGAG” sequence that is also located at the 3’-terminus of the 60bp-6 substrate (Figure 3A, marked with yellow). This finding provides a plausible explanation for both the retained binding observed with the 60bp-7-V substrate and the weak but detectable binding previously noted with the 60bp-6 substrate. These results collectively indicate that GCG6mAG may not represent the minimal recognition motif of VchI.

**Figure 2.**
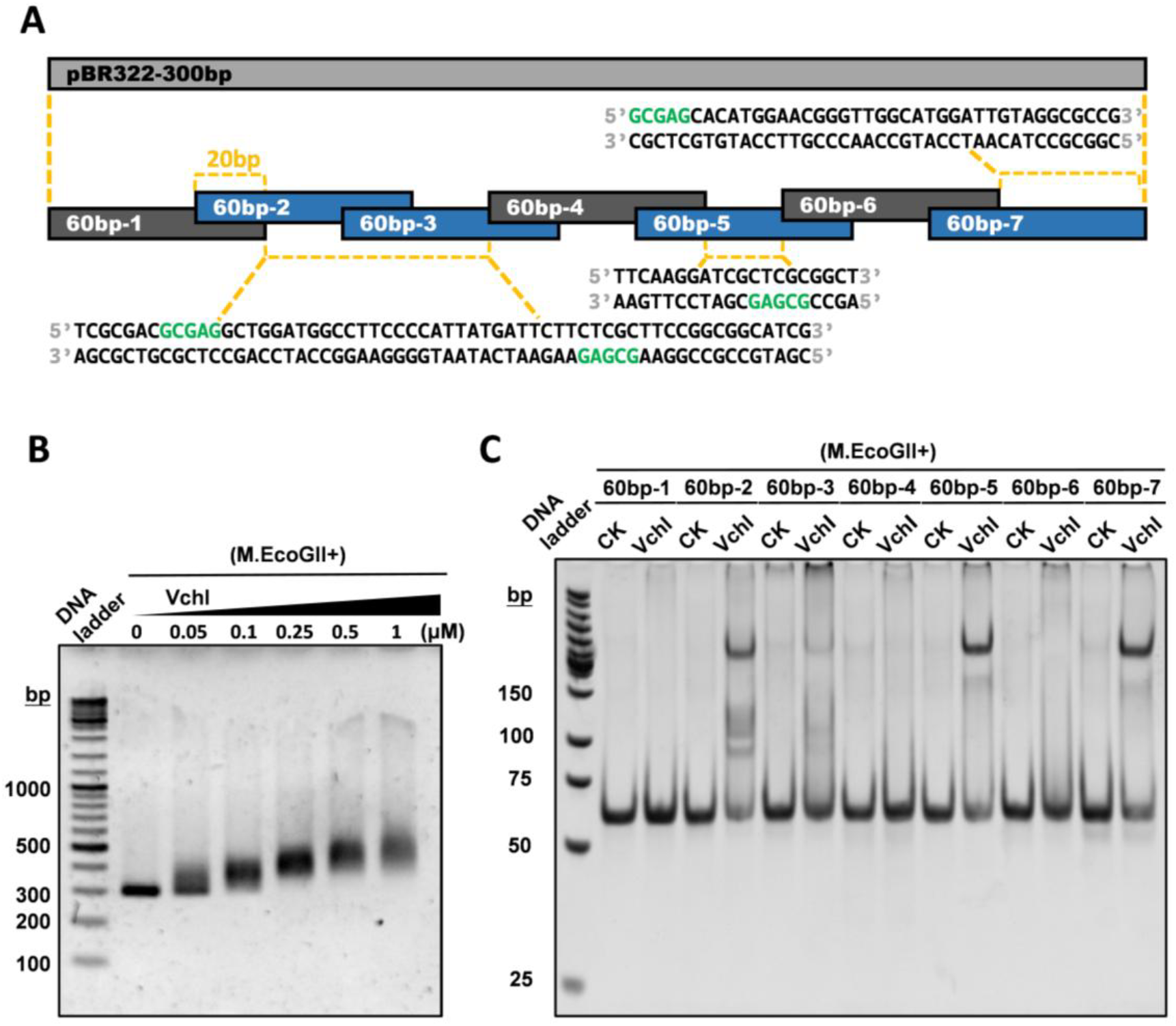
Characterization of binding motifs. **(A)** Schematic diagram of the 300 bp DNA substrate sequence segmentation. In the lower part of this figure, the colored sequences are the consensus sequences of the binding fragments. Specifically, red indicates N6 - methyladenine (6mA), and green indicates unmodified bases. **(B)** The binding activity of VchI at varying concentrations to a 300 bp DNA fragment methylated by M.EcoGII was analyzed by EMSA. The specific primers are shown in Table S2. **(C)** The binding activity of VchI to seven 60 bp fragments methylated by M.EcoGII was analyzed by EMSA, with “CK” indicating the negative control without VchI. The specific information of the sequences is shown in Table S3.

**Figure 3.**
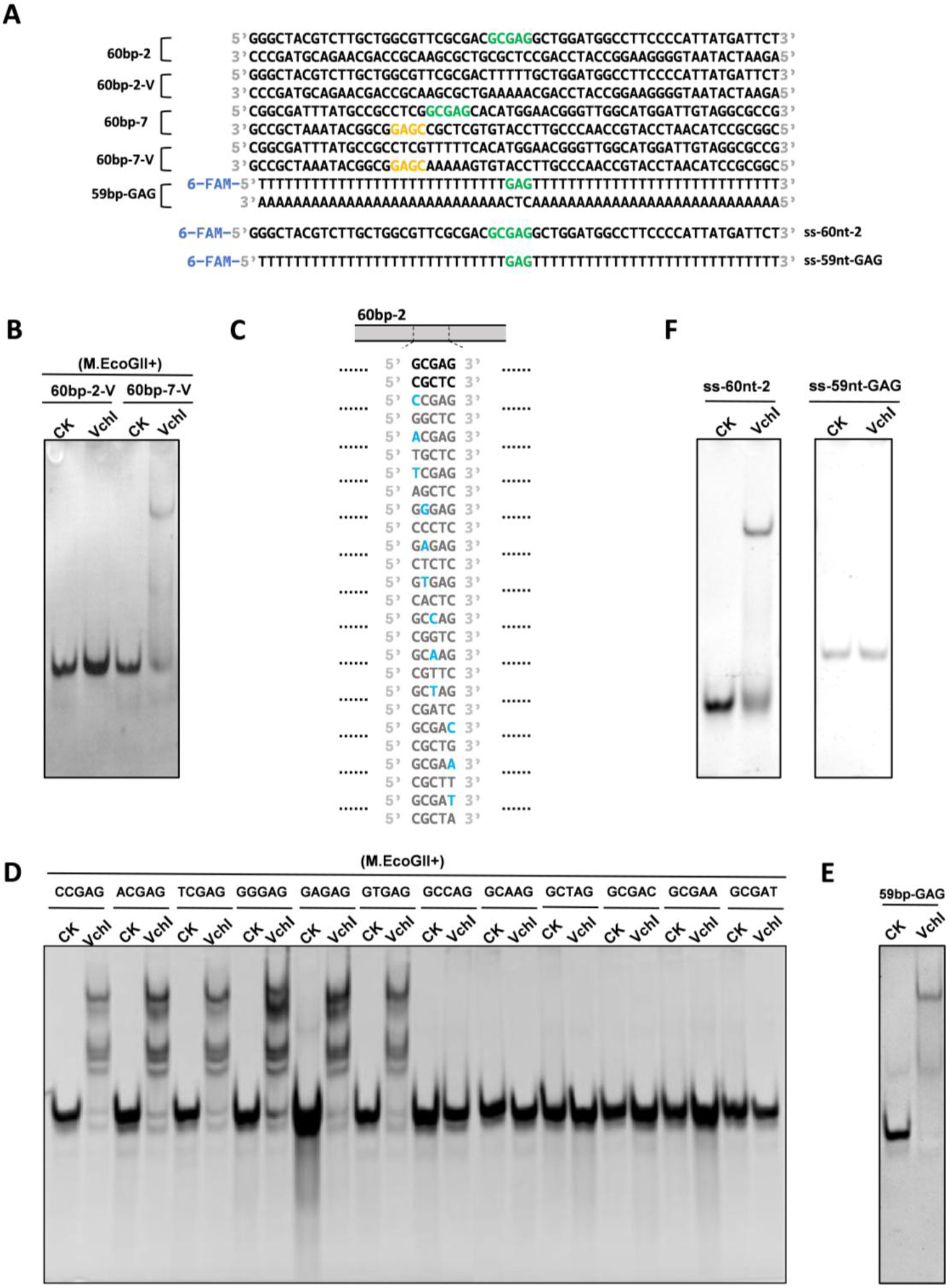
In-depth characterization of binding motifs. **(A)** Schematic representation of the substrates used in **(B, E, F)**. The 60bp-2 and 60bp-7 sequences and their corresponding mutants, and synthetic sequences (59bp-GAG, ss-60nt-2 and ss-59nt-GAG), with color-highlighted motifs indicate the binding sequence regions. **(B)** The binding of VchI to M.EcoGII-methylated substrates (60 bp-2-V and 60 bp-7-V) was analyzed by EMSA. **(C)** Schematic representation of the substrates used in **(D)**. Twelve single-nucleotide variants of the 60bp GCGAG sequence, where blue-labeled bases denote the substituted nucleotides. **(D)** The binding of VchI to twelve M. EcoGII-methylated mutants was analyzed by EMSA. **(E)** The binding of VchI to M.EcoGII-methylated substrate (59nt-GAG) was analyzed by EMSA. **(F)** The binding of VchI to M.EcoGII-methylated substrates (ss-60nt-2 and ss-59nt-GAG) was analyzed by EMSA. The specific information of the sequences is shown in Table S3. *The VchI concentrations used in all the above experiments were consistently maintained at 0.5 μM. “CK” represents the negative control without VchI.

We performed comprehensive single-nucleotide mutagenesis in the “GCGAG” sequence within the 60bp-2 fragment to further characterize the core of the VchI recognition motif. Twelve mutant substrates were generated by substituting each non-adenine position with all three alternative nucleotides (Figure 3C). Following *in vitro* methylation using M. EcoGII, each modified substrate was incubated with VchI and analyzed by EMSA (Figure 3D). Mutations at the first (G) and second (C) positions showed minimal impact on VchI binding affinity. In contrast, substitutions at either the third (G) or fifth (G) positions resulted in complete loss of binding capacity to VchI. These results demonstrate that VchI specifically recognizes the G6mAG motif. To demonstrate that the G6mAG motif alone is sufficient for VchI binding, we synthesized a 59-bp DNA substrate containing only the G6mAG motif flanked by A-T base pairs. EMSA confirmed that VchI exhibited robust binding to this substrate (Figure 3E), providing conclusive evidence that the G6mAG motif represents the essential and sufficient recognition element for VchI. We further tested whether VchI recognizes the G6mAG motif in single-stranded (ss) DNA. A 60-nt ssDNA corresponding to the G6mAG-containing strand of 60bp-2 (ss-60nt-2) and a 59-nt ssDNA corresponding to the G6mAG-containing strand of 59bp-G6mAG (ss-59nt-G6mAG) were incubated with VchI respectively. EMSA showed that VchI exhibited a certain degree of binding to the ss-60nt-2 substrate but no binding to the ss-59nt-G6mAG substrate (Figure 3F). These findings suggest that VchI’s binding to the G6mAG-containing DNA depends on a double-stranded (ds) DNA conformation. The observed binding between VchI and the ss-60nt-2 substrate may result from its potential secondary structure that mimics local dsDNA.

## Discussion

Although the wH domain of VchI is homologous to those in DpnI and HHPV4I (Figure 4A), two key residues involved in 6mA recognition differ between them (V42 and R45 in VchI; K229 and Q232 in DpnI; K197 and Q200 in HHPV4I) (Figure 4B). Considering that both DpnI and HHPV4I recognize the G6mATC motif whereas VchI recognizes the G6mAG motif, these residues may contribute to the discrepancy in the recognition specificities. It was demonstrated that mutations at these residues (V42K and R45A in VchI) completely abolished the binding capacity of VchI to G6mAG-containing DNA (Figure 4C, D), confirming their critical role in G6mAG recognition. These findings advance our understanding of the 6mA recognition mechanism of the wH domain and provide insights for engineering 6mA-binding proteins with customized recognition motifs.

**Figure 4.**
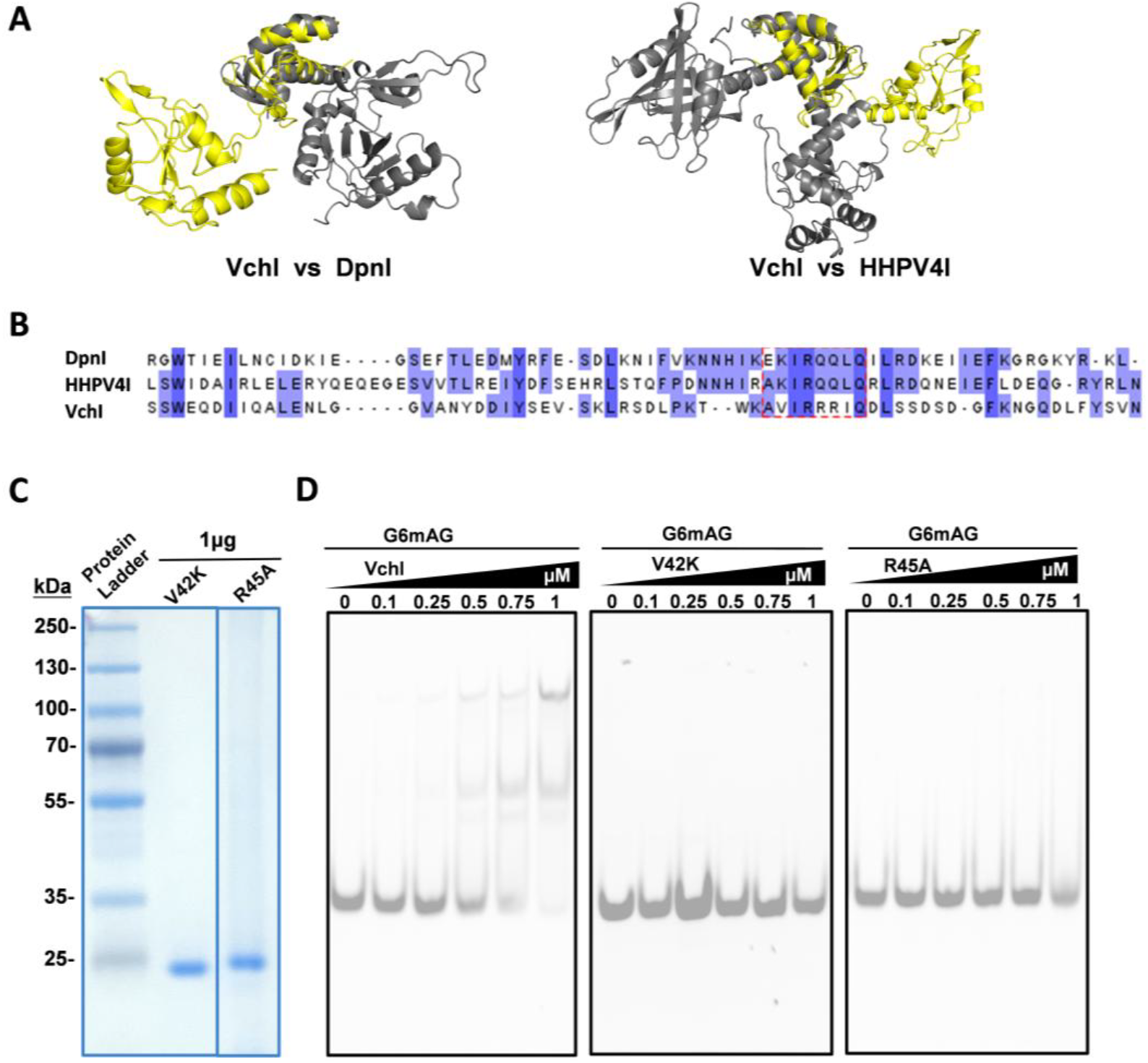
Detection of the wH domain mutants of VchI. **(A)**. The three-dimensional structural comparison of VchI (yellow) with DpnI and HHPV4I (gray) in the wH domain. **(B)**. Potential key residues in the wH domain were identified based on sequence alignment of the wH domains of DpnI, HHPV4I and VchI. **(C)** SDS-PAGE analysis of the purified VchI mutants (V42K and R45A). **(D)** Electrophoretic mobility shift assay (EMSA) was performed M.EcoGII-methylated 19-bp dsDNA substrate incubated with increasing concentrations of VchI or its mutants (V42K and R45A). The specific sequence of the 19-bp dsDNA is shown in Table S3.

Notably, despite harboring an HNH endonuclease domain, VchI displayed no detectable DNA cleavage activity across various tested conditions including different metal ions, pH levels, and salt concentrations (Supplementary Figure S1). We hypothesized that VchI may function as a DNA-binding protein involved in epigenetic regulation, rather than serving as a component of a conventional restriction-modification system.

## Acknowledgements

We thank all lab members for helpful discussions. This work was supported by the National Natural Science Foundation of China (grant 32150009), the Feng Foundation of Biomedical Research, and the Billionhome Venture Capital.

