## Supplemental Figure1, Supplemental Table for "A Gm6AG-binding protein from *Vibrio cholerae*"

This file contains Supplementary Figures S1 and Supplementary Tables S1-S3.

### Figure S1

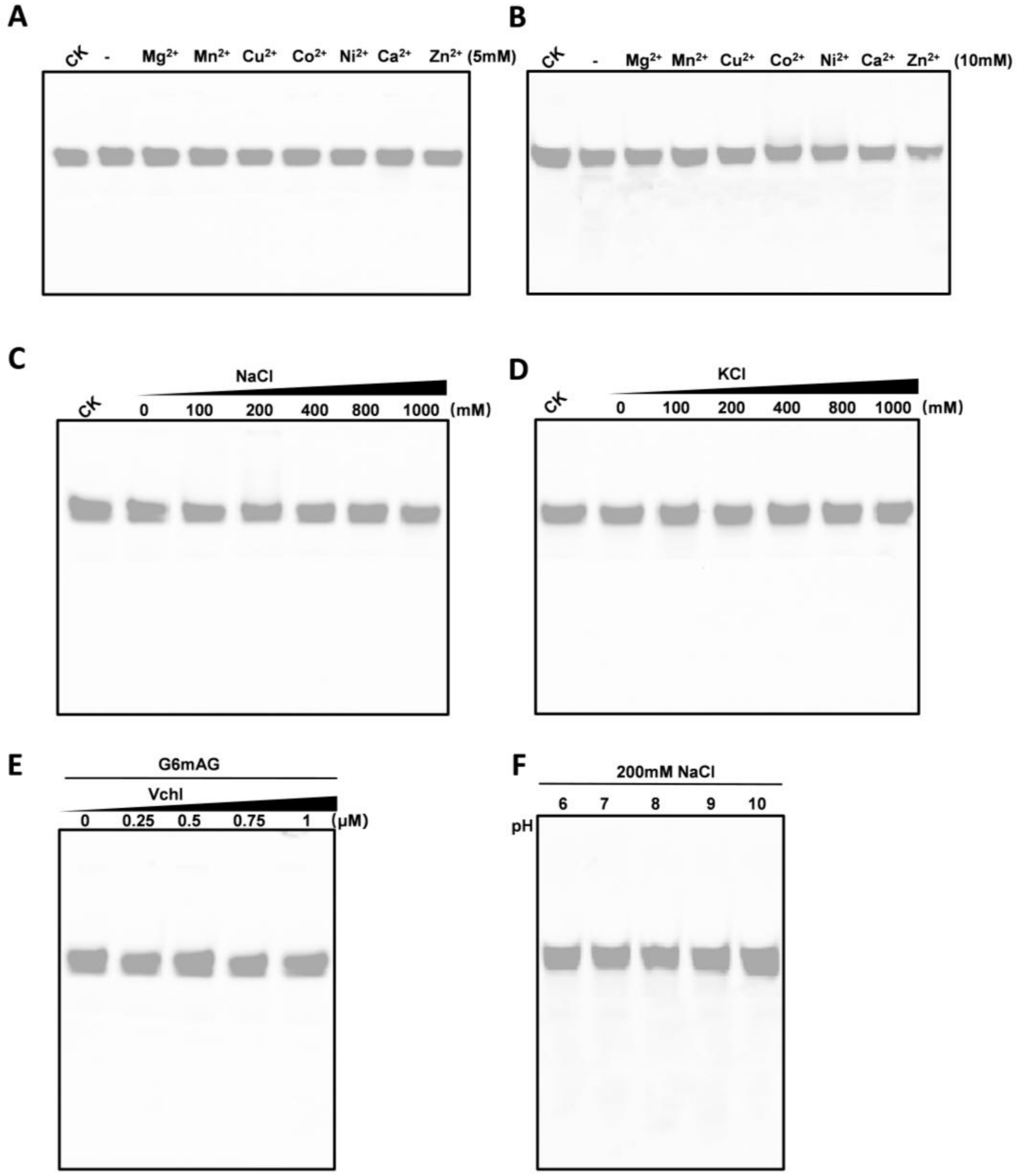


**Figure 6. Cleavage activity of VchI.**

**(A and B)** The dsDNA cleavage activity of VchI in different divalent metal ions. The reaction system containing 50 mM Tris-HCl (pH 7.9), 100 mM NaCl and 5 mM of specified different types of metal ions with 0.5 μM VchI and 100 ng of 19bp dsDNA. **(C and D)** The DNA cleavage activity of VchI in the presence of different concentrations of NaCl and KCl. Different concentrations of NaCl or KCl (100, 200, 400, 800, 1000 mM) were added to the enzymatic digestion reaction system with 0.5 μM VchI and 100 ng of 19bp dsDNA. **(E)** The dsDNA cleavage activity of VchI at different concentrations. Incubate varying concentrations (0.25, 0.5, 0.75, 1 μM) of VchI with 100 ng of 19bp dsDNA in the restriction digestion reaction system. **(F)** DNA cleavage activity of VchI at different pH**.** React 0.5 μM VchI with 100 ng of 19bp dsDNA in restriction digestion buffers at different pH values (6.0, 7.0, 8.0, 9.0, 10.0). *The symbol "-" indicates that no metal ions are added to the system, and "CK" represents the blank control without adding VchI in the system. All reactions were performed in 10 μL volume at 37°C for 1 hour.

Table S1. Plasmids and primers for protein expression.

| Sequence/Primer name | sequence（5’-3’） |
|---|---|
| Optimized VchI gene sequence 633 bp | ATGAGTTCATGGGAACAGGATATTATACAAGCGCTGGAAAACCTGGGCG<br>GTGTTGCGAATTATGATGATATTTATTCTGAAGTCTCCAAACTGCGTAGC<br>GACTTGCCAAAGACCTGGAAAGCTGTGATTCGTCGCCGTATCCAGGATCT<br>CAGTTCTGACAGCGACGGCTTCAAGAACGGTCAGGACTTGTTTTACAGCG<br>TTAATGGCTTGGGCGGTGGCATGTGGGGTCTGCGTAATAAGTTAGCCTAC<br>ACTCCGAAGGCGGCAGATCTGCCGACCGGTACAAAAGAGCCGGAACGTG<br>AGTATACGACCACCTATCGTGTTTTGCGCGATACCAATCTGGCGCGTAAA<br>CTTAAGTTGCTGTACAACAACTCCTGCCAGATCTGCGGTCTGCAAATCCA<br>ACTGCCGAACGGCAAATTGTACTCTGAAGCGCACCACATCATCCCGCTGG<br>GTAATCCGCATCATGGTAGCGACACCCCGGAGAACATTATCGTGCTGTGT<br>CCGAACCACCACGTGATGTGCGACTACGGCGCAATTGCTCTCTCGCTGAA<br>GGAGGTGAAGCAGGTTAGCTCCCACAGCATCAGCCAGAAAAGCATTGAT<br>TACCATAACAAAATCATCCGCGAGACGGAACTGTAA |
| VchI WP_057555222.1 210 aa | MSSWEQDIIQALENLGGVANYDDIYSEVSKLRSDLPKTWKAVIRRRIQDLSSD<br>SDGFKNGQDLFYSVNGLGGGMWGLRNKLAYTPKAADLPTGTKEPEREYTTT<br>YRVLRDTNLARKLKLLYNNSCQICGLQIQLPNGKLYSEAHHIIPLGNPHHGSD<br>TPENIIVLCPNHHVMCDYGAIALSLKEVKQVSSHSISQKSIDYHNKIIRETEL |
| pQE80L-F | TAATTAGCTGAGCTTGGACTCCTGT |
| pQE80L-R | ATGGTGATGGTGATGGTGAGATCCT |
| VchI-F | TCACCATCACCATCACCATATGAGTTCATGGGAACAGGATATTATAC |
| VchI-R | GTCCAAGCTCAGCTAATTATTACAGTTCCGTCTCG |
| V42A-F | GGAAAGCTGCGATTCGTCGCCGTATCCAG |
| V42A-R | CGACGAATCGCAGCTTTCCAGGTCTTTGG |
| R44A-F | CTGTGATTGCGCGCCGTATCCAGGATC |
| R44A-R | TGGATACGCGCACGAATCACAGCTTTCCAGGTC |
| R45A-F | GTGATTCGTGCCCGTATCCAGGATCTCAGTTC |
| R45A-R | TGGATACGGGCACGAATCACAGCTTTCCAG |
| pET28a-F | GAATTCGAGCTCCGTCGACAAGCTTG |
| pET28a-R | CATATGGCTGCCGCGCGGCACCAGG |

Table S2. Primers for Fragments used in Figure 1.

| Primer name | sequence（5’-3’） |
|---|---|
| T7-3 kb-F | CTCTTTCGTTACGTGAACGAATCCGTGAGCACCTA |
| T7-3 kb-R | TTAAACACAACATGTTCAACTGGGGTGTAAGGAG |

| | |
|---|---|
| pBR-2385 bp-F | GCAGGCCATGCTGTCCAGGCAGGTAGATGACGACCAT |
| pBR-2385 bp-R | AAGTTGGGTGCACGAGTGGGTTACATCGAACTGGATCT |
| pBR-300 bp-F | AGAAGCAGGCCATTATCGCCGGCAT |
| pBR-300 bp-R | CGGCGCCTACAATCCATGCCAACCC |

Table S3. Fragments used in Figure 2, 3,4, S1.

| Fragment name | sequence（5’-3’） |
|---|---|
| 60 bp-1-F | AGAAGCAGGCCATTATCGCCGGCATGGCGGCCGACGCGCTGGGCTAC GTCTTGCTGGCGT |
| 60 bp-1-R | ACGCCAGCAAGACGTAGCCCAGCGCGTCGGCCGCCATGCCGGCGATA ATGGCCTGCTTCT |
| 60 bp-2-F | GGGCTACGTCTTGCTGGCGTTCGCGACGCGAGGCTGGATGGCCTTCCC CATTATGATTCT |
| 60 bp-2-R | AGAATCATAATGGGGAAGGCCATCCAGCCTCGCGTCGCGAACGCCAG CAAGACGTAGCCC |
| 60 bp-3-F | GCCTTCCCCATTATGATTCTTCTCGCTTCCGGCGGCATCGGGATGCCC GCGTTGCAGGCC |
| 60 bp-3-R | GGCCTGCAACGCGGGCATCCCGATGCCGCCGGAAGCGAGAAGAATCA TAATGGGGAAGGC |
| 60 bp-4-F | GGATGCCCGCGTTGCAGGCCATGCTGTCCAGGCAGGTAGATGACGAC CATCAGGGACAGC |
| 60 bp-4-R | GCTGTCCCTGATGGTCGTCATCTACCTGCCTGGACAGCATGGCCTGCA ACGCGGGCATCC |
| 60 bp-5-F | TGACGACCATCAGGGACAGCTTCAAGGATCGCTCGCGGCTCTTACCA GCCTAACTTCGAT |
| 60 bp-5-R | ATCGAAGTTAGGCTGGTAAGAGCCGCGAGCGATCCTTGAAGCTGTCC CTGATGGTCGTCA |
| 60 bp-6-F | CTTACCAGCCTAACTTCGATCACTGGACCGCTGATCGTCACGGCGATT TATGCCGCCTCG |
| 60 bp-6-R | CGAGGCGGCATAAATCGCCGTGACGATCAGCGGTCCAGTGATCGAAG TTAGGCTGGTAAG |
| 60 bp-7-F | CGGCGATTTATGCCGCCTCGGCGAGCACATGGAACGGGTTGGCATGG ATTGTAGGCGCCG |
| 60 bp-2-cut-F | GGGCTACGTCTTGCTGGCGTTCGCGACTTTTTGCTGGATGGCCTTCCCC ATTATGATTCT |
| 60 bp-2-cut-R | AGAATCATAATGGGGAAGGCCATCCAGCAAAAAGTCGCGAACGCCAG CAAGACGTAGCCC |
| 60 bp-7-cut-F | CGGCGATTTATGCCGCCTCGTTTTTCACATGGAACGGGTTGGCATGGA TTGTAGGCGCCG |
| 60 bp-7-cut-R | CGGCGCCTACAATCCATGCCAACCCGTTCCATGTGAAAAACGAGGCG GCATAAATCGCCG |

| | |
|---|---|
| 59 bp-GAG-F | TTTTTTTTTTTTTTTTTTTTTTTTTTTTG$_{M6}$AGTTTTTTTTTTTTTTTTTTTT TTTTTTTT |
| 59 bp-GAG-R | AAAAAAAAAAAAAAAAAAAAAAAAAAAACTCAAAAAAAAAAAAAA AAAAAAAAAAAAAA |
| CCGAG-F | GGGCTACGTCTTGCTGGCGTTCGCGACCCGAGGCTGGATGGCCTTCCC CATTATGATTCT |
| CCGAG-R | AGAATCATAATGGGGAAGGCCATCCAGCCTCGGGTCGCGAACGCCAG CAAGACGTAGCCC |
| ACGAG-F | GGGCTACGTCTTGCTGGCGTTCGCGACACGAGGCTGGATGGCCTTCCC CATTATGATTCT |
| ACGAG-R | AGAATCATAATGGGGAAGGCCATCCAGCCTCGTAAAAAAAAAAAAA AAAAAAAAAAAAAA |
| TCGAG-F | GGGCTACGTCTTGCTGGCGTTCGCGACTCGAGGCTGGATGGCCTTCCC CATTATGATTCT |
| TCGAG-R | AGAATCATAATGGGGAAGGCCATCCAGCCTCGAGTCGCGAACGCCAG CAAGACGTAGCCC |
| GGGAG-F | GGGCTACGTCTTGCTGGCGTTCGCGACGGGAGGCTGGATGGCCTTCCC CATTATGATTCT |
| GGGAG-R | AGAATCATAATGGGGAAGGCCATCCAGCCTCCCGTCGCGAACGCCAG CAAGACGTAGCCC |
| GAGAG-F | GGGCTACGTCTTGCTGGCGTTCGCGACGAGAGGCTGGATGGCCTTCCC CATTATGATTCT |
| GAGAG-R | AGAATCATAATGGGGAAGGCCATCCAGCCTCTCGTCGCGAACGCCAG CAAGACGTAGCCC |
| GTGAG-F | GGGCTACGTCTTGCTGGCGTTCGCGACGTGAGGCTGGATGGCCTTCCC CATTATGATTCT |
| GTGAG-R | AGAATCATAATGGGGAAGGCCATCCAGCCTCACGTCGCGAACGCCAG CAAGACGTAGCCC |
| GCCAG-F | GGGCTACGTCTTGCTGGCGTTCGCGACGCCAGGCTGGATGGCCTTCCC CATTATGATTCT |
| GCCAG-R | AGAATCATAATGGGGAAGGCCATCCAGCCTGGCGTCGCGAACGCCAG CAAGACGTAGCCC |
| GCAAG-F | GGGCTACGTCTTGCTGGCGTTCGCGACGCAAGGCTGGATGGCCTTCCC CATTATGATTCT |
| GCAAG-R | AGAATCATAATGGGGAAGGCCATCCAGCCTTGCGTCGCGAACGCCAG CAAGACGTAGCCC |
| GCTAG-F | GGGCTACGTCTTGCTGGCGTTCGCGACGCTAGGCTGGATGGCCTTCCC CATTATGATTCT |
| GCTAG-R | AGAATCATAATGGGGAAGGCCATCCAGCCTAGCGTCGCGAACGCCAG CAAGACGTAGCCC |
| GCGAC-F | GGGCTACGTCTTGCTGGCGTTCGCGACGCGACGCTGGATGGCCTTCCC CATTATGATTCT |
| GCGAC-R | AGAATCATAATGGGGAAGGCCATCCAGCGTCGCGTCGCGAACGCCAG CAAGACGTAGCCC |

| | |
|---|---|
| GCGAA-F | GGGCTACGTCTTGCTGGCGTTCGCGACGCGAAGCTGGATGGCCTTCCC CATTATGATTCT |
| GCGAA-R | AGAATCATAATGGGGAAGGCCATCCAGCTTCGCGTCGCGAACGCCAG CAAGACGTAGCCC |
| GCGAT-F | GGGCTACGTCTTGCTGGCGTTCGCGACGCGATGCTGGATGGCCTTCCC CATTATGATTCT |
| GCGAT-R | AGAATCATAATGGGGAAGGCCATCCAGCATCGCGTCGCGAACGCCAG CAAGACGTAGCCC |
| 19 bp-F | CGCGACGCG6mAGGCTGGATG |
| 19 bp-R | CATCCAGCCTCGCGTCGCG |
